# Multimodal gene embeddings for drug-target prediction and lineage reconstruction

**DOI:** 10.64898/2026.02.13.689089

**Authors:** Benjamin L. Kidder

## Abstract

Understanding how gene function emerges across molecular, cellular, and pharmacologic contexts remains a central challenge in systems biology and drug discovery. Conventional computational models typically operate within a single modality, such as expression, ontology, or interaction networks, limiting their ability to capture the multidimensional nature of gene function. Here, we present NEWT (Neural Embeddings for Wide-spectrum Targeting), a multimodal deep learning framework that integrates heterogeneous biological knowledge into a unified and interpretable representation space. By combining functional annotations, large-scale co-expression data, pathway information, lineage programs, transcriptional regulons, and protein-protein interaction features through an attention-guided fusion architecture, NEWT learns cross-modal dependencies that reflect both global functional hierarchies and context-specific regulatory relationships. Applied to L1000 perturbational transcriptomes, NEWT achieves higher compound-target prediction accuracy than prior embedding models and reconstructs pharmacological networks that reveal mechanistic and repurposing opportunities. When extended to single-cell RNA-seq data, NEWT preserves developmental trajectories and enhances the resolution of lineage hierarchies. Together, these results demonstrate that multimodal gene embeddings can bridge pharmacogenomic and single-cell transcriptomic analyses within a common functional geometry, establishing a scalable foundation for integrative target discovery and systems-level modeling of cellular identity.

## INTRODUCTION

The rapid growth of transcriptomic and functional genomics data has transformed how gene function and cellular phenotypes are analyzed. High-throughput platforms such as RNA-seq, the Connectivity Map (CMap), and large-scale perturbation studies now enable systematic mapping of compound-gene-cell relationships across diverse contexts^1–3^. Despite recent progress, most computational approaches remain limited in the breadth of modalities they integrate, capturing only a subset of the molecular, regulatory, and functional dimensions that define gene activity. This constraint reduces both predictive accuracy and interpretability across molecular and cellular scales, particularly for applications such as drug repurposing, target prioritization, and lineage reconstruction.

Drug development pipelines remain slow and costly, with high attrition rates in late-stage trials^4^. Computational models that can accurately infer drug-target associations from molecular data offer a route to accelerate therapeutic discovery and repositioning. Network-based inference^5, 6^, graph embeddings^7, 8^, and deep learning approaches^9, 10^ have shown promise in integrating heterogeneous molecular data. Among these, gene-centric embeddings, continuous vector representations that encode functional relationships, have become a powerful paradigm for representing complex biological systems. For example, Gene2Vec^11^ and OPA2Vec^12^ learn distributed representations from gene co-expression or ontology contexts, while clusDCA^13^ uses random walks on Gene Ontology graphs. These models capture aspects of gene co-functionality, but they typically rely on a single information source and lack mechanisms for context-specific integration across modalities.

Recent frameworks have extended these ideas toward compound-target prediction by embedding transcriptional perturbation signatures. Notably, FRoGS^14^ demonstrated that deep embeddings of L1000 gene expression profiles can improve drug-target recall compared to conventional similarity metrics. However, FRoGS and similar architectures operate primarily on expression and ontology information, without explicit integration of regulatory or network-level context. As a result, such models may overlook complementary biological information that could enhance predictive performance and generalization across datasets.

In parallel, single-cell RNA-sequencing (scRNA-seq) has revolutionized our understanding of cellular diversity and developmental hierarchies^15–17^. Computational advances now allow millions of single cells to be profiled across tissues and perturbations, revealing dynamic trajectories and lineage transitions. Despite these breakthroughs, most single-cell methods embed cells rather than genes, and are therefore optimized for clustering or visualization rather than transfer learning of functional gene relationships. Integrating gene-level embeddings that encode regulatory and network knowledge into single-cell analysis could bridge the gap between molecular perturbations and cell-state identity.

Despite progress, current gene embedding strategies remain fragmented in scope and application. Most models are modality-specific, trained on co-expression, ontology, or perturbation data alone, and fail to capture the composite functional landscape of genes across cellular contexts and regulatory hierarchies. Their performance often depends strongly on the source modality and the downstream task, reflecting limited cross-domain generalizability. Moreover, embedding frameworks developed for drug-target prediction rarely incorporate single-cell information or account for lineage and state relationships, constraining their ability to transfer gene-level representations to cell-level inference.

Conversely, existing single-cell embedding methods typically model cells as independent entities without leveraging gene-centric embeddings that encode molecular function and network connectivity. A unified framework is therefore needed, one that integrates multimodal functional genomics data, transfers these representations to perturbational and pharmacogenomic contexts, and applies them to single-cell analyses to reveal lineage structure and cell-state transitions within a common embedding space.

To address these limitations, we developed NEWT (Neural Embeddings for Wide-spectrum Targeting), an independently implemented and modular framework for multimodal gene embedding and cross-omics prediction. NEWT integrates functional annotations from Gene Ontology^18^, large-scale co-expression data from ARCHS4^19^, curated pathway signatures from MSigDB^20^, lineage-specific regulatory networks from CellNet^21^, transcription-factor regulons from DoRothEA^22^ and CollecTRI^23^, and protein-protein interaction embeddings^24^. These heterogeneous data sources are combined through an attention-guided fusion architecture that learns modality-specific weighting, enabling context-aware gene representations that generalize across tasks.

Applied to L1000 perturbational transcriptomes, NEWT achieves improved recall and precision in compound-target association relative to single-modality and FRoGS-based models^14^. The model captures cross-modal dependencies that reflect complementary aspects of gene function, such as regulatory modules that link co-expression clusters with protein interaction neighborhoods. When extended to single-cell datasets such as PBMC3k^25^, the same multimodal embeddings preserve developmental topology and enhance clustering resolution, outperforming baseline PCA and single-source embeddings in delineating hematopoietic lineages. These findings suggest that multimodal gene embeddings can simultaneously serve as a unifying representation for pharmacogenomic modeling and as a biologically interpretable coordinate system for single-cell state analysis.

Beyond performance gains, NEWT introduces a modular and reproducible workflow accessible through command-line utilities for model training, visualization, and ATC-based network export. The framework is implemented in Python with TensorFlow and Scanpy back ends and supports transparent benchmarking across data modalities and biological tasks. By unifying pharmacogenomic and single-cell transcriptomic analyses within a shared embedding space, NEWT facilitates data-driven hypothesis generation and integrative exploration of how molecular perturbations shape cellular phenotypes.

In summary, this study presents a multimodal deep learning framework that integrates diverse biological data sources to learn gene representations consistent across molecular and cellular contexts. The resulting embeddings improve drug-target prediction, preserve lineage architecture in single-cell analyses, and establish a scalable foundation for integrative functional genomics. The framework is openly available at: https://github.com/KidderLab/NEWT.

## RESULTS

### Multimodal gene embeddings reveal cohesive and biologically grounded organization of tissue-specific programs

Representing gene function in a way that reflects biological context remains a central challenge in functional genomics. Previous approaches, such as those embedding gene signatures into functional spaces derived from Gene Ontology (GO) and ARCHS4 co-expression data^14^, have improved compound-target prediction but remain limited in scope. These representations rely primarily on ontological and expression-based information, without integrating complementary biological priors such as regulatory networks, pathway context, or protein-protein interactions. To overcome these constraints, we developed a multimodal attention-based framework that learns unified gene representations by jointly integrating diverse sources of biological knowledge.

Each gene was embedded across complementary knowledge sources encompassing Gene Ontology, ARCHS4 co-expression networks, MSigDB pathway annotations, CellNet lineage networks, transcription factor regulons from DoRothEA and CollecTRI, and protein-protein interaction (PPI) graphs. These modality-specific embeddings were then fused through a neural attention mechanism that learns the relative contribution of each data source for tissue-specific gene classification^26–28^. The performance of each embedding combination is summarized in **Table S1**, which includes all concatenation-based and fusion-based models.

Performance benchmarking across all modality combinations demonstrated that multimodal integration improved accuracy relative to individual modalities, although fusion did not uniformly exceed the strongest concatenation-based models. The highest overall performance was achieved by the PPI + CellNet combination (accuracy 0.9833). The strongest fusion model, Default + CellNet + CollecTRI, performed nearly identically (accuracy 0.9831). Several other multimodal combinations incorporating lineage information and regulatory features also performed strongly, whereas single-modality models based on DoRothEA, CollecTRI, MSigDB, or PPI alone showed markedly lower accuracy. These findings, detailed in **Table S1**, highlight the dominant influence of lineage-level structure on tissue classification and the added value of complementary knowledge sources.

When visualized in two dimensions, multimodal embeddings generated by full-data concatenation displayed clear separation between the three tissue groups, with each lineage forming a coherent region of the projection despite some expected internal variability (**Fig. 1A**). Genes originating from the same tissue aggregated closely, whereas those from different tissues were separated robustly, reflecting biologically meaningful relationships encoded in the combined representation. In comparison, the GO + ARCHS4 baseline^14^ (**Fig. 1B**) produced broader and partially overlapping clusters, and the one-hot identity control (**Fig. 1C**) showed no discernible structure.

**Figure 1.**
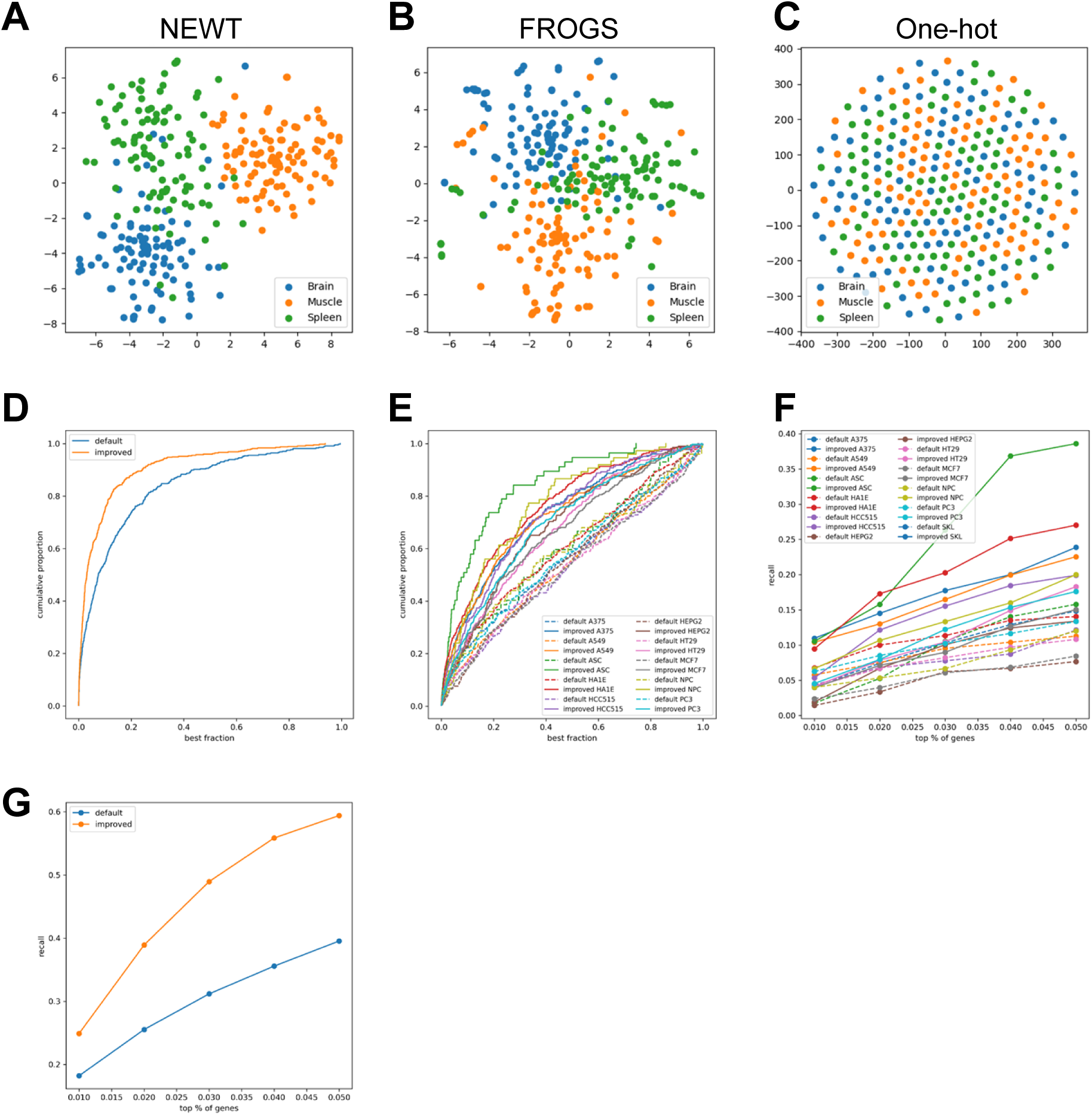
Multimodal fusion captures biologically grounded organization of gene function. (A) Two-dimensional projection of fused multimodal gene embeddings showing tissue-specific organization of gene representations. Each point represents a gene colored by its dominant tissue of expression, forming distinct, non-overlapping clusters that correspond to coherent biological programs. (B) FRoGS-default embedding (GO + ARCHS4) displaying broader, partially overlapping tissue distributions, indicating reduced resolution compared with NEWT. (C) One-hot identity baseline showing random dispersion of genes without discernible biological structure. (D) Classification accuracy across ten cross-validation folds comparing multimodal, FRoGS-default, and one-hot models. (E) Precision-recall and receiver-operating-characteristic (ROC) curves illustrating superior performance of the multimodal model. (F) Ranking of model performance across all single-and multi-modality configurations. (G) Cumulative accuracy distribution across modality combinations showing consistent dominance of multimodal fusion models.

### Integration of complementary biological sources enhances discriminative power

We next assessed whether the structural improvements of the multimodal representation corresponded to better tissue separability. Across the full range of cumulative classification thresholds, NEWT exceeded the performance of the FROGS baseline, indicating more discriminative organization of tissue-specific signatures (**Fig. 1D**). Precision-recall and ROC analyses further demonstrated uniformly higher curves for NEWT across tissues (**Fig. 1E**). These results were supported by direct cross-validation accuracy estimates, in which multimodal combinations incorporating CellNet and additional regulatory modalities achieved the strongest performance, including accuracies of 0.9833 for the PPI + CellNet model and 0.9831 for the Default + CellNet + CollecTRI fusion model.

### Robust performance and high top-k recall across heterogeneous contexts

Ranking analyses provided a comprehensive view of performance across all single-and multimodal combinations. When ordered by test accuracy, models enriched for lineage-informed features such as CellNet consistently occupied the highest positions, regardless of whether the features were combined through direct concatenation or through the attention-based fusion mechanism (**Fig. 1F**). Configurations dominated by CellNet, with or without PPI or additional regulatory sources, formed a clear cluster at the top of the ranking, whereas weak single-modality models clustered at the bottom.

The cumulative performance distribution further illustrated these trends (**Fig. 1G**). Multimodal combinations spanned the upper accuracy range and showed a smoother and more favorable cumulative profile compared with the default approach, which accumulated rapidly in the lower accuracy region. Together, these results demonstrate that integrating complementary biological priors, particularly lineage-level and regulatory context, produces stronger and more stable predictions, reflecting a more coherent organization of functional relationships within the embedding space.

### Multimodal embeddings redefine cellular architecture and reveal emergent lineage logic

To examine whether multimodal integration can reorganize the structure of single-cell transcriptomic data, we applied the NEWT framework to the well-characterized PBMC3k dataset^29^ (10x Genomics; Scanpy tutorial dataset) encompassing major immune populations. Unlike embeddings derived solely from transcriptomic or ontological features, NEWT unifies orthogonal biological priors, including ontological hierarchies, co-expression dynamics, pathway associations, transcription factor regulons, and protein-protein interaction networks, within a shared representational space. In this space, genes are arranged not merely by expression correlation but by their contextual role within regulatory and structural networks. This integration yields a geometry in which transcriptomic variation is interpreted through biological relationships rather than statistical proximity, providing a more meaningful substrate for single-cell inference. For this analysis, we used the NEWT multimodal concatenated embedding rather than the attention-based fusion model, because the joint PCA + NEWT representation provides stable and well-structured geometry for single-cell visualization.

When visualized in two dimensions, the NEWT joint embedding produced an ordered cellular landscape (**Fig. 2A-C**). The PCA projection (**Fig. 2A**) separated major populations but lacked fine-grained organization within or between compartments. In contrast, the GO + ARCHS4 baseline embedding (**Fig. 2B**) generated a diffuse manifold with incomplete delineation of lymphoid and myeloid lineages. The NEWT multimodal concatenated embedding (**Fig. 2C**), integrated with the top principal components, resolved sharply defined and non-overlapping clusters corresponding to canonical immune lineages while also revealing substructure that was not apparent in either PCA or ontology-based spaces. B-cell and plasma cell populations localized adjacently but remained distinct, whereas NK, T-cell, and myeloid compartments occupied separated regions arranged in a pattern consistent with developmental and functional proximity. The resulting topology recapitulates the latent geometry of hematopoietic differentiation, capturing continuity where lineage relationships exist and orthogonality where divergence has occurred, demonstrating that multimodal integration restructures single-cell variation into a biologically coherent spatial organization.

**Figure 2.**
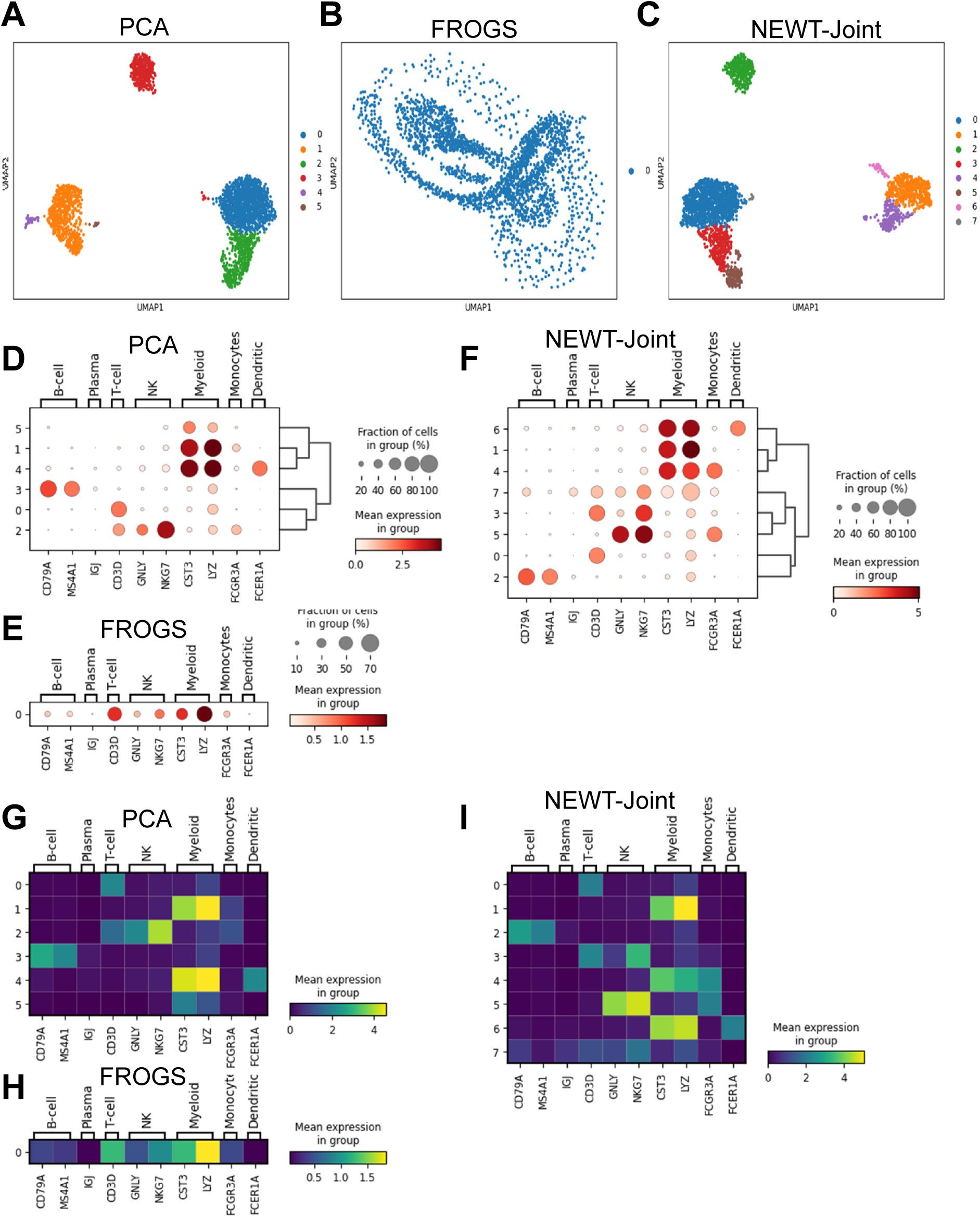
Multimodal embeddings reconstruct cellular hierarchy and lineage organization in single-cell data. (A) PCA projection of PBMC3k single-cell RNA-seq data showing separation of major immune lineages. (B) FRoGS-default (GO + ARCHS4) embedding showing diffuse cluster. (C) NEWT joint multimodal embedding displaying sharply defined, non-overlapping clusters corresponding to immune lineages. Monocytes and dendritic populations are distinctly resolved. (D-F) Dotplots of canonical lineage marker expression confirming cluster identity. PCA (D) shows broadly distributed marker expression, FRoGS-default (E) shows loss of resolution, and the NEWT-joint embedding (F) exhibits confined domains corresponding to lineage boundaries. (G-I) Matrixplots showing mean marker expression per cluster. PCA (G) exhibits moderate clustering but limited lineage definition, FRoGS-default (H) shows reduced resolution, and the NEWT-joint embedding (I) displays clear block-diagonal modularity consistent with lineage-specific regulatory coordination.

Within the myeloid compartment, the NEWT joint embedding sharpened the delineation of classical (LYZ⁺) and non-classical (FCGR3A⁺) monocytes and enhanced the definition of the dendritic cell cluster marked by FCER1A expression (**Fig. 2C, F, I**). These subsets appeared conflated or poorly resolved in the PCA and GO + ARCHS4 embeddings, where related myeloid populations formed partially overlapping groups. The distinct separation of these closely related lineages reflects regulatory and signaling differences captured by the multimodal priors, demonstrating that NEWT preserves fine-grained sub-lineage organization that conventional transcriptomic embeddings portray with reduced resolution.

Examination of canonical lineage markers confirmed that NEWT isolates transcriptional identities with a clarity unattainable in lower-order embeddings (**Fig. 2D-F**). B-cell markers (CD79A, MS4A1) were confined to a single contiguous cluster, whereas CD3D defined discrete T-cell islands, and cytotoxic genes (GNLY, NKG7) localized exclusively to NK cells. Myeloid markers (LYZ, CST3, FCGR3A) and dendritic signatures (FCER1A) formed cohesive, non-overlapping modules. In contrast, these signatures were broadly distributed across clusters in the PCA and GO + ARCHS4 embeddings, underscoring that multimodal integration distinguishes lineage-specific transcriptional variance from background co-expression. The joint embedding therefore resolves transitional or mixed states without fragmenting legitimate cell types.

Expression heatmaps and matrixplots (**Fig. 2G-I**) further demonstrated that NEWT imposes a modular structure aligned with regulatory coordination rather than purely co-expression. Within the same analytical framework, NEWT produces diagnostic visualizations that trace these relationships across scales, linking modular architecture to lineage continuity and marker specificity.

**Supplementary Fig. S1A** shows the global UMAP of the NEWT joint embedding colored by Leiden clusters. **Supplementary Fig. S1B-C** demonstrate that canonical lineage markers and lineage-associated gene modules align coherently with these clusters.

**Supplementary Fig. S2** compared marker-level organization across four embedding strategies. The FROGS default embedding (**Supplementary Fig. S2A**) failed to recover any discernible structure, producing a uniform heatmap with no visible alignment of lineage markers. The multimodal-only NEWT embedding (**Supplementary Fig. S2B**) introduced partial organization, with early grouping of related markers but incomplete separation of lineage modules. While PCA alone (**Supplementary Fig. S2C**) captured variation, the NEWT joint embedding that integrates PCA with the multimodal representation (**Supplementary Fig. S2D**) further refined this organization into a modular and hierarchical pattern, with lineage-specific genes forming well defined modules. These comparisons indicate that multimodal biological priors alone provide limited improvements, whereas coupling them with transcriptomic variance through the joint embedding yields the most coherent and interpretable representation of cellular organization.

Gene-level trackplots (**Supplementary Fig. S3**) revealed clear transitions in CD79A, CD3D, and LYZ expression across neighboring clusters, indicating that the NEWT joint embedding preserves transcriptional continuity and arranges related immune states in a manner consistent with established lineage relationships. Expression heatmaps overlays (**Supplementary Fig. S4**) showed that lineage markers were more sharply segregated and spatially coherent in the NEWT joint embedding than in PCA alone.

Marker-wise violin plots (**Supplementary Fig. S5**) showed that lineage-specific genes such as CD79A and MS4A1 displayed sharply confined, unimodal expression within the expected clusters in the NEWT joint embedding. In contrast, the default FROGS embedding showed no meaningful cluster structure, and both PCA and GO + ARCHS4 embeddings produced broader or partially multimodal distributions. The NEWT joint model exhibited the most refined and coherent clustering, with clearer delineation of lineage-restricted expression compared with PCA alone. Quality-control metrics, including n_genes and percent_mito, were similar across all embeddings (**Supplementary Fig. S6**), confirming that the improved resolution arises from representational strength rather than technical bias.

Collectively, these findings demonstrate that the NEWT joint embedding reconstructs cellular organization with high structural and functional fidelity. By positioning gene activity within a biologically informed coordinate system, NEWT converts diffuse single-cell representations into coherent manifolds that reveal discrete lineage identities and preserve the continuity of related immune states. The resulting embedding integrates molecular, regulatory, and network-derived priors to generate an interpretable view of cell identity that aligns expression variation with established lineage structure. In doing so, NEWT provides a general framework for representing cellular diversity in single-cell datasets.

### Network-level organization and functional modularity of multimodal compound-target predictions

Having established that NEWT embeddings capture coherent transcriptional and lineage structure at single-cell resolution, we next evaluated how this representational framework extends to compound-target inference and network organization. By applying multimodal integration to perturbational transcriptomes, NEWT embeds small-molecule responses within the same biologically informed coordinate system, enabling direct alignment of compound-induced signatures with their mechanistic gene targets and associated pathways.

To examine how multimodal functional embeddings reorganize compound–target relationships, we applied NEWT to shRNA-derived perturbational profiles from the Library of Integrated Network-Based Cellular Signatures (LINCS) L1000 dataset^2^. This analysis generated a global compound-target network in which nodes correspond to genes and edges denote predicted associations derived from similarity in the multimodal embedding space. In concept, this extends the embedding-based framework introduced previously^14^, but NEWT incorporates a broader set of biological priors, resulting in a representation that improves the modular organization and interpretability of compound-target connectivity.

The global shRNA-derived compound-target network inferred by NEWT (**Fig. 3A**) formed an extensive and interconnected graph in which compounds and their predicted gene targets grouped into coherent regions that reflected ATC class relationships and shared functional programs. Targets linked to several ATC categories acted as connectors between major regions, while genes with more restricted pathway roles were positioned toward the periphery. ATC labels were used only for downstream visualization and were not provided to the model during training.

**Figure 3.**
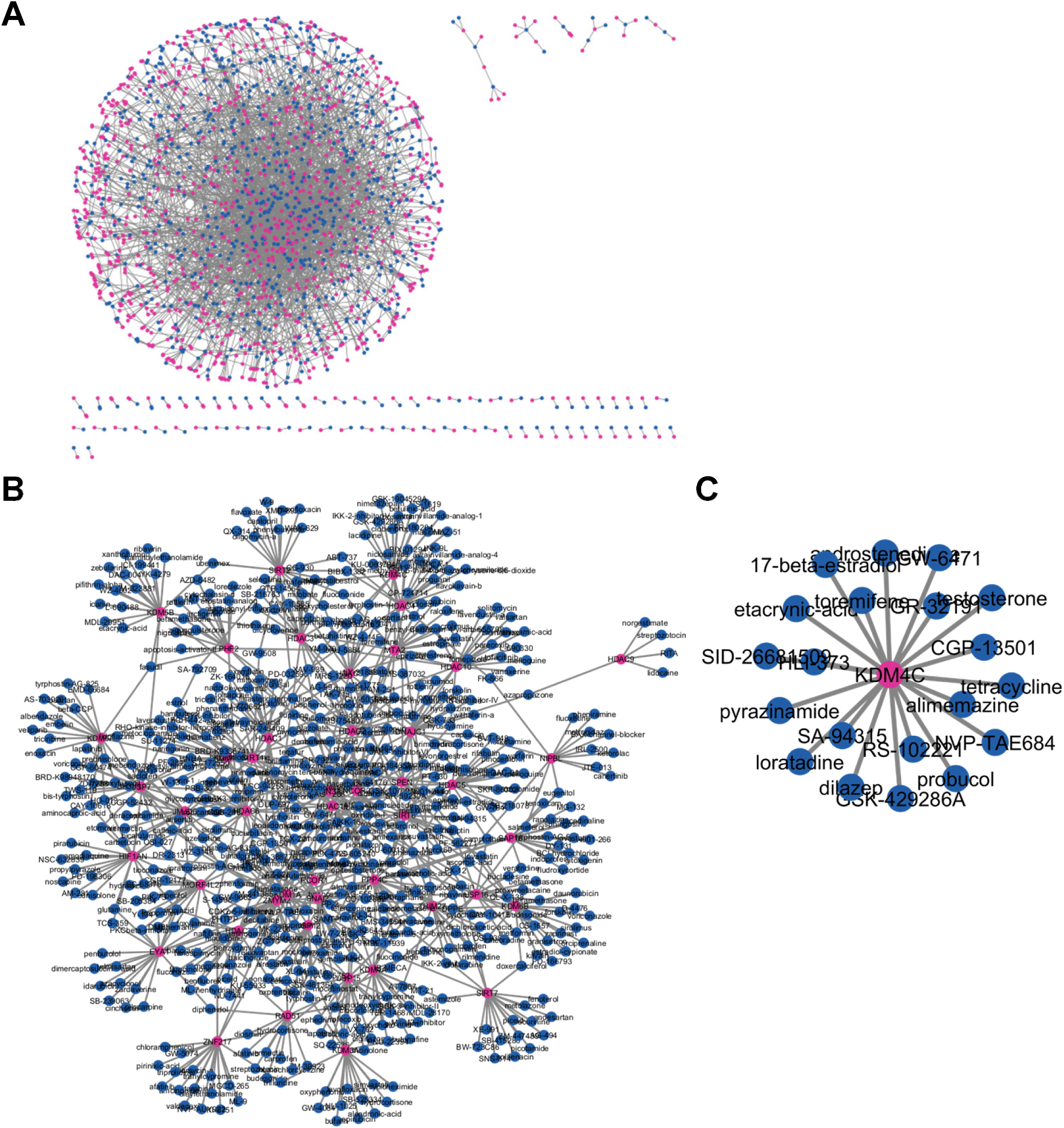
Multimodal network analysis reveals pharmacologically coherent compound-target modules. (A) Global compound-target network derived from NEWT predictions using shRNA perturbation profiles. Nodes represent genes (magenta) or compounds (blue), and edges denote predicted associations. The graph exhibits a modular topology with a densely connected regulatory core and peripheral subnetworks corresponding to distinct biological processes. (B) Epigenetic subnetworks (EpiGenes) extracted from the same global network, showing connectivity among chromatin regulators and RNA-modifying enzymes. Central hubs link multiple compound classes, reflecting integrative control across chromatin and signaling pathways. (C) Representative subnetwork highlighting epigenetic regulator and associated compounds predicted to target histone demethylation, transcriptional elongation, and RNA modification processes.

Summary statistics for the global network are reported in **Table S3**. The final graph contained 5289 nodes (1211 compounds and 4078 targets) and 276582 unique edges. The network showed mean degrees of 228 for compounds and 68 for targets, along with a median target degree of 54. Although the overall density was low (0.0198), the graph displayed clear internal structure, with five communities detected and a modularity score of 0.109. These properties indicate a globally dense network that still organizes along major functional axes consistent with the transcriptional and regulatory programs captured by NEWT.

Within the epigenetic gene subset, chromatin-associated regulators were used to extract focused subnetworks from the global graph. The resulting EpiGenes networks (**Fig. 3B-C**) displayed highly connected regions where transcriptional regulators, chromatin remodelers, and DNA or histone modification enzymes converged with compounds predicted to influence them. These structures remained stable despite the density of the global graph, indicating that epigenetic regulators form reproducible functional neighborhoods within the multimodal NEWT representation.

Prominent regulators such as BRD4, KDM4C, and DNMT1 occupied central positions, linking compounds across multiple mechanistic categories. Their placements reflect the broad influence of chromatin accessibility, methylation control, and transcriptional regulation on compound-induced expression changes. These hubs also acted as connectors between regions that would otherwise form separate partitions.

Community analysis within the EpiGenes subnetworks identified groups that correspond to known chromatin-associated activities, including histone demethylation, chromatin remodeling complexes, transcriptional elongation machinery, and RNA-modifying enzymes. Genes participating in related regulatory pathways displayed shared compound interactions, indicating that NEWT organizes epigenetic regulators according to coordinated regulatory logic rather than isolated gene-specific responses. Subclusters enriched for RNA-modification enzymes and chromatin remodelers were located adjacently, consistent with relationships between transcriptional control and post-transcriptional regulation.

Together, these network structures show that multimodal integration preserves established biological relationships while also organizing pharmacological space into coherent modules that reveal mechanistic connections between druggable regulators. By embedding compound-induced expression responses within a biologically informed coordinate system, NEWT provides a framework for identifying related targets, guiding repurposing, and uncovering previously unrecognized regulatory associations.

### Multimodal embeddings reveal a structured landscape of biological processes

To evaluate how NEWT organizes biological function across the genome, we visualized the fused multimodal gene embeddings using two-dimensional t-SNE projections (**Fig. 4A-B**). Each point represents a gene embedded within a joint representation space constructed from ontology, co-expression, pathway, regulatory, and interaction priors. Although t-SNE preserves primarily local relationships, the fused NEWT embedding itself encodes broader structure, and this underlying organization is reflected in the low-dimensional projections. Genes associated with transcriptional regulation, signaling, immune activation, and cell-cycle control occupied distinct yet related regions, while metabolic and structural pathways formed continuous gradients between adjacent functional neighborhoods.

**Figure 4.**
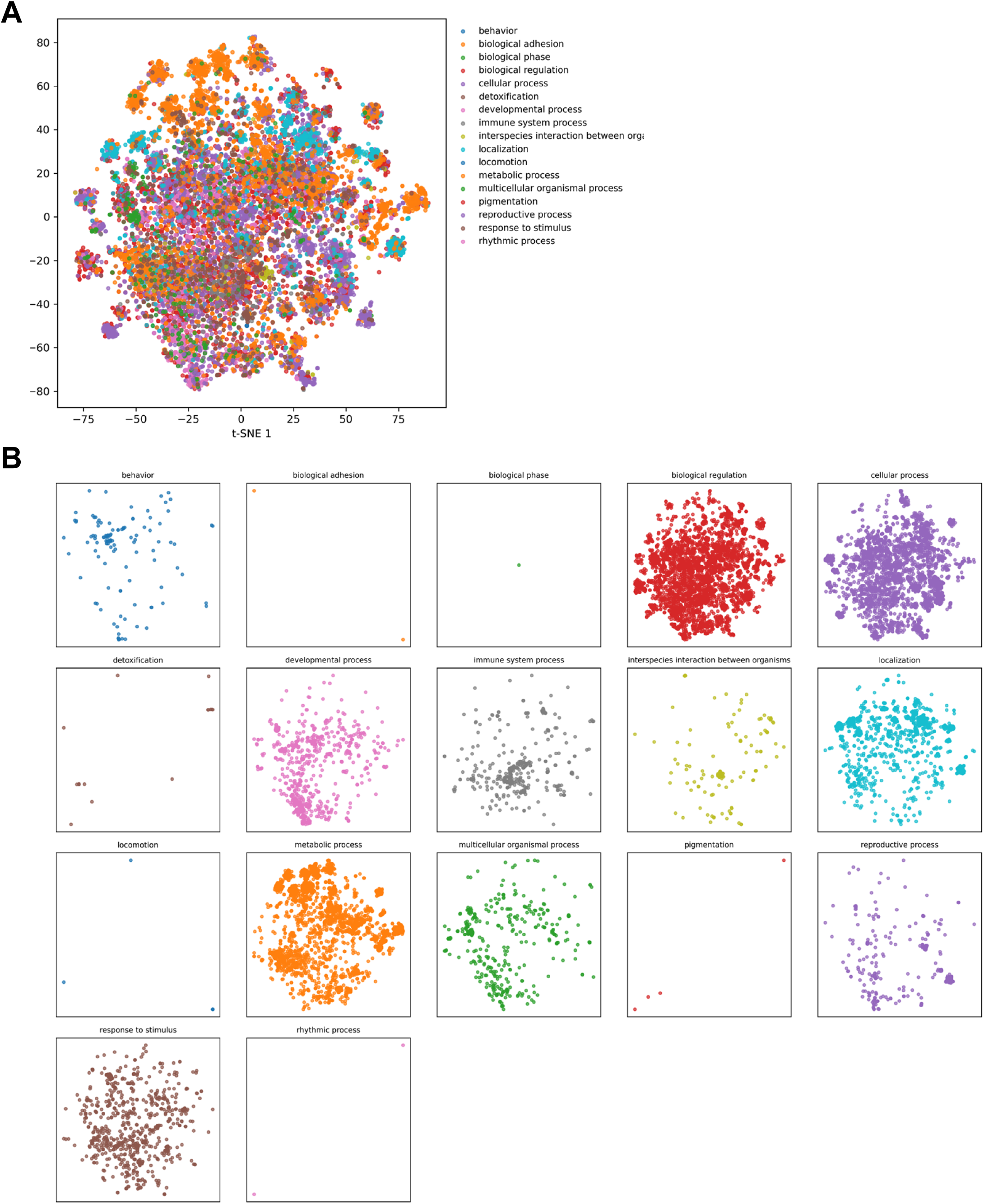
Multimodal gene embeddings reveal the global organization of biological processes. (A) Two-dimensional t-SNE embedding of NEWT multimodal gene representations colored by top-level Gene Ontology (GO) process. Genes cluster by biological domain, including transcriptional regulation, signaling, immune response, metabolism, and development. (B) Per-category mosaic visualization showing process-specific t-SNE maps. Compact clusters for biological regulation, cellular process, and metabolic process highlight the modular organization of the fused embedding, while related processes such as chromatin organization and RNA metabolism appear in adjacent regions linked by shared regulatory annotations. Together, these projections show that NEWT embeddings capture the hierarchical and interconnected architecture of cellular function within a unified coordinate system for gene-level biological relationships.

Category-level mosaic visualizations (**Fig. 4B**) showed that most top-level Gene Ontology processes were spatially coherent within the fused embedding. Processes such as biological regulation, cellular process, and metabolic process formed compact regions, indicating that NEWT captures the modular architecture of biological systems. Related functional groups, including immune response, signal transduction, and developmental processes, appeared as neighboring regions linked through shared regulatory and transcriptional factors, consistent with functional coupling between adaptive signaling and differentiation.

Continuity between conceptually related categories was also evident. Chromatin organization, transcriptional regulation, and RNA metabolic processes formed a connected axis, reflecting the coupling between regulatory and transcriptional programs preserved by multimodal integration.

Together, these analyses show that NEWT generates a biologically interpretable functional manifold in which local and global patterns arise from the combination of complementary biological priors. The fused embedding unifies ontological, regulatory, and interaction information into a continuous coordinate system that captures discrete pathway boundaries as well as the transitional relationships that connect them. NEWT therefore provides a scalable framework for functional discovery across molecular and systems contexts.

## DISCUSSION

This study introduces NEWT (Neural Embeddings for Wide-spectrum Targeting), a multimodal deep learning framework that integrates heterogeneous sources of biological knowledge into a unified representation space for mechanistic target prediction and inference of cellular states.

By integrating ontological structure^18^, co-expression patterns^19^, pathway context^20^, lineage programs^21^, transcription-factor regulons^22, 23^, and protein-interaction networks^24^, NEWT captures functional dependencies that extend beyond any individual modality. Across applications such as compound-target prediction and single-cell lineage reconstruction, the unified embedding revealed emergent biological organization and demonstrated improved mechanistic interpretability and structural coherence compared with prior single-source or bimodal approaches^2, 14^.

### A multimodal representation of gene function

The conceptual strength of NEWT lies in framing gene representation as a problem of multimodal fusion rather than feature concatenation. Its attention-guided architecture dynamically weighs input modalities, learning context-dependent relationships between ontology, co-expression, and regulatory information^26–28^. This approach contrasts with frameworks such as Gene2Vec^11^ and OPA2Vec^12^, which consider individual knowledge sources independently. Mechanistically, the model integrates lineage-level structure from CellNet^21^ with transcriptional specificity from DoRothEA and CollecTRI^22, 23^. The resulting embeddings form a continuous functional manifold in which genes self-organize according to shared biological logic.

### Bridging pharmacogenomics and network medicine

Projecting L1000 perturbational profiles^2^ into the fused space enabled NEWT to reconstruct a global compound-target network informed by biological semantics. The resulting hierarchy includes central hubs that connect pharmacologic classes across chromatin regulation, metabolism, and transcriptional control, consistent with principles of network medicine^5^ and with broad regulatory influence on transcriptional responses. Although the global graph is dense, it contains interpretable communities that highlight functional coherence within chromatin remodeling, RNA modification, histone demethylation, and transcriptional elongation. These features extend earlier embedding-based frameworks^14^ and demonstrate that multimodal integration improves intra-module structure and cross-module interpretability.

The context-awareness of NEWT opens avenues for rational drug repurposing. Compounds positioned within the same multimodal neighborhood share underlying regulatory architecture rather than relying solely on expression-level similarity. This representation therefore complements efforts to map pharmacogenomic interactions^3^ and to apply machine learning to drug discovery pipelines^10^. Within the epigenetic landscape, EpiGenes subnetworks revealed convergent regulatory organization, consistent with established relationships between chromatin remodeling and transcriptional progression. Such mechanistic clusters may guide the identification of new targets or repositioning of existing agents within therapeutically relevant pathways.

### Cellular topology and lineage fidelity

Applied to single-cell RNA-seq data^25, 29^, NEWT embeddings recapitulated canonical immune hierarchies while resolving fine-grained distinctions such as FCGR3A-positive monocytes. This behavior aligns with integrative single-cell frameworks^16^ yet differs in a key aspect: NEWT embeds genes rather than cells, enabling a unified coordinate system that generalizes across bulk perturbational data and cellular state space. The geometry of the multimodal manifold reflects both continuity and boundary. Developmental trajectories appear as smooth transitions between adjacent modules, whereas distinct cell states maintain separation. These properties parallel the continuity described in hematopoietic epigenomics^17^ and show that multimodal embeddings can recover lineage logic directly from expression data without explicit trajectory models.

### Mechanistic and conceptual implications

NEWT shows that gene function can be represented as an emergent property of overlapping regulatory, ontological, and interaction hierarchies. Prior work has shown that deep embedding models can capture biological semantics across multiple modalities^10, 26^. The fused embedding acts as a coordinate system in which genes arrange themselves according to shared regulatory logic, co-expression, and physical interaction context. This reflects systems-level principles of modular organization^5, 6^, where biological behavior arises from coordinated activity within interconnected subnetworks. Within NEWT, this modularity arises without explicit supervision on pathways, indicating that the model has internalized latent structure present across functional genomics datasets.

By embedding diverse priors into a unified representation, NEWT moves beyond correlation-based similarity networks and toward context-aware functional inference, where proximity encodes shared mechanism rather than mere statistical association. Mechanistically, the performance of NEWT across compound-target prediction and single-cell analysis shows that multimodal fusion yields a transferable scaffold that generalizes across biological scales. The same manifold that organizes pharmacologic targets by regulatory class also orders immune and developmental lineages by transcriptional hierarchy.

### Translational outlook

The translational value of NEWT lies in its capacity to unify discovery and interpretation across domains. In pharmacology, multimodal embeddings can support target identification, anticipate off-target interactions, and enable mechanism-driven compound repurposing. In systems and developmental biology, NEWT provides a representation for understanding how regulatory networks govern lineage transitions, informing precision strategies in regenerative medicine. In data science, the architecture offers a scalable foundation for integrating heterogeneous omics datasets, aligning with the trajectory of emerging attention-based multi-omics models^26, 27^.

## Conclusion

NEWT provides an attention-guided representation of gene function that integrates diverse biological priors into a coherent and interpretable structure. By bridging perturbational genomics and single-cell transcriptomics, the framework unifies compound-target prediction and cell-state inference within a shared embedding space. This integration enables mechanistic generalization across scales and contexts, transforming multidimensional biological data into operational structure. As multimodal embeddings expand to incorporate new experimental and clinical data types, frameworks such as NEWT may translate systems-level insights into predictive, experimentally testable hypotheses that accelerate therapeutic development and advance precision medicine.

## MATERIALS & METHODS

### Framework overview

NEWT (Neural Embeddings for Wide-spectrum Targeting) was developed as a multimodal deep learning framework that unifies heterogeneous sources of biological information into shared, context-aware gene representations. The workflow integrates ontology, co-expression, pathway, regulatory, lineage, and interaction-derived features into a neural architecture with attention-guided fusion. This multimodal design enables consistent mapping of genes, compounds, and single cells into a common functional coordinate system, allowing inference across molecular, pharmacologic, and transcriptomic domains.

The analysis pipeline consisted of three major stages: (i) generation of modality-specific embeddings and attention-fused multimodal representations; (ii) application of the learned embeddings to compound-target prediction and network reconstruction using perturbational profiles; and (iii) transfer of these embeddings into single-cell RNA-seq data to test whether the fused functional geometry reconstructs cellular hierarchies. The framework was implemented in Python using TensorFlow for neural components and Scanpy for single-cell workflows.

### Data sources and preprocessing

NEWT embeddings were trained on complementary biological data modalities representing distinct facets of gene function.

#### Ontology context

Gene Ontology^18^ (GO; release 2024-03) provided the hierarchical structure of biological processes, molecular functions, and cellular components. Binary term membership matrices were filtered to exclude overly broad (>2,000 genes) or sparse (<10 genes) categories.

#### Co-expression dynamics

Large-scale RNA-seq co-expression data were obtained from ARCHS4^19^. Pairwise gene-gene correlations were computed and projected into a 256-dimensional principal component space to capture dominant transcriptional covariation patterns across tissues.

#### Pathway associations

Pathway-level membership vectors were derived from the Molecular Signatures Database (MSigDB^20^).

#### Lineage programs

CellNet^21^ provided tissue-and lineage-specific regulatory network connectivity scores. Each gene’s connectivity to lineage modules was normalized to form a continuous vector representing its tissue association.

#### Regulatory priors

Transcription-factor regulon data were integrated from DoRothEA^22^ and CollecTRI^23^, providing directionality and confidence-weighted TF-target interactions.

#### Protein interaction topology

Protein-protein interaction data were obtained from STRING^30^, which provides confidence-weighted human interaction networks. A graph-based embedding was derived from these interactions to capture both local neighborhood structure and broader network topology.

All sources were harmonized to Entrez Gene identifiers and aligned to a common reference universe of protein-coding genes. Missing values were imputed as zeros, redundant entries were averaged, and all modalities were normalized to zero mean and unit variance prior to model training.

### Multimodal embedding architecture and model training

Each modality was processed independently through a two-layer feedforward encoder using rectified linear unit activations and layer normalization. Encoders were optimized to reconstruct modality-specific features, ensuring that information within each biological domain was preserved prior to fusion. The encoded representations were concatenated and passed to a self-attention layer that learned to assign dynamic weights to each modality based on its relevance to the training objective. This attention-guided fusion mechanism produced a single 256-dimensional embedding for each gene.

Training proceeded in two phases. During pretraining, a self-supervised reconstruction objective aligned the latent spaces of all modalities. During fine-tuning, the model was trained to classify tissue-specific gene signatures provided in a curated label file derived from CellNet lineage and tissue annotations. Optimization employed the Adam optimizer with a learning rate of 1 × 10⁻³, cosine annealing, dropout (0.3), and early stopping based on validation loss.

To assess reproducibility and generalization, ten-fold cross-validation was performed on the gene-level lineage classification task, and model performance was benchmarked against (i) single-modality encoders, (ii) the FRoGS-default model using GO and ARCHS4 embeddings^14^, and (iii) one-hot encodings. Performance metrics included accuracy, precision, recall, F1-score, and AUROC.

Attention weights were extracted and averaged across folds to determine the relative importance of each data source, revealing which modalities contributed most to the learned functional geometry. These embeddings formed the foundation for downstream analyses.

### Compound-target prediction and network analysis

To evaluate whether multimodal embeddings could improve pharmacologic inference, compound-target predictions were generated from perturbational gene-expression data derived from the Library of Integrated Network-Based Cellular Signatures (LINCS) L1000 dataset^2^. For each compound, differential expression profiles were correlated with those produced by shRNA-mediated gene knockdowns across multiple cell types. Connectivity scores between compounds and target genes were computed and used to train a fully connected neural classifier in which the NEWT gene embeddings served as features replacing raw differential expression values as input features. Known drug-target relationships from DrugBank^31^ and ChEMBL^32^ were used as positive supervision for binary classification of predicted associations with negative examples sampled from gene-compound pairs lacking known interactions.

Predicted compound-target associations were ranked by confidence and used to construct a weighted bipartite graph in which nodes represented genes or compounds and edges corresponded to predicted association strength. Graph metrics including node degree, betweenness centrality, and modularity were computed using NetworkX. Communities were detected using the Louvain algorithm, and functional enrichment was assessed using g:Profiler and Enrichr.

To focus on epigenetic regulation, subnetworks enriched for chromatin remodelers, histone demethylases, and RNA modification enzymes were extracted from the global compound-target graph, forming the EpiGenes subnetworks. These subnetworks were visualized in Cytoscape^33^ and analyzed for inter-module connectivity, revealing hierarchical organization among pharmacologically tractable chromatin regulators.

### Single-cell RNA-seq integration

To test whether the learned multimodal gene embeddings generalize to cellular contexts, the model was applied to the 10x Genomics PBMC3k single-cell RNA-seq dataset^29^. After standard filtering (cells with <500 genes or>10% mitochondrial reads removed), expression counts were log-normalized and scaled to unit variance.

Each gene in the expression matrix was replaced with its corresponding NEWT embedding vector, and each cell’s representation was computed as the expression-weighted average of its constituent gene embeddings with genes lacking embeddings ignored or assigned zero vectors to maintain dimensional consistency. This approach effectively projects cells into the same multimodal coordinate space learned from bulk functional data, allowing direct visualization of cellular organization within the integrated functional manifold.

Dimensionality reduction was performed using t-distributed stochastic neighbor embedding (t-SNE) and UMAP. Clusters were identified using the Leiden algorithm, and marker-based cell type annotation was conducted using canonical immune lineage genes.

Comparisons were made to PCA-and FRoGS-default-based embeddings to assess improvements in cluster separability and lineage continuity. Visual inspection of UMAP projections and marker-gene expression confirmed that the multimodal embedding preserved developmental structure and enhanced lineage resolution across immune populations.

### Visualization of multimodal landscapes

The global organization of biological processes was visualized using t-SNE applied to the final fused embedding matrix rather than to individual modalities. Each point corresponded to a gene, and colors reflected its top-level Gene Ontology category. Additional t-SNE projections were generated for each GO term, and results were assembled into mosaic visualizations summarizing the distribution of major functional processes.

Distances between GO category centroids were computed using Euclidean metrics to infer higher-order proximity among biological processes. Functional adjacency networks were generated to visualize the modular continuity between transcriptional regulation, signaling, metabolism, and developmental pathways. These visualizations formed the basis of Figure 4, demonstrating how multimodal fusion reconstructs the latent topology of biological function.

## DECLARATIONS

### ETHICS STATEMENT

All analyses were based on de-identified, publicly available datasets.

### CONSENT FOR PUBLICATION

All authors have read and approved the final version of this manuscript.

### Statistical Analysis

Quantitative performance comparisons were based on accuracy, precision-recall, and receiver operating characteristic metrics computed using scikit-learn. Reported values represent mean ± standard deviation across cross-validation folds.

### Data and Code Availability

All datasets used in this study are publicly available. Gene Ontology and ARCHS4 co-expression embeddings were obtained from the FRoGS repository^14^ (https://github.com/chenhcs/FRoGS) and used as baseline modalities for multimodal fusion.

Pathway information was sourced from MSigDB (https://www.gsea-msigdb.org).

Regulatory priors were downloaded from CellNet (https://cellnet.hms.harvard.edu), DoRothEA (https://saezlab.github.io/dorothea), and CollecTRI (https://saezlab.github.io/collectri).

Protein-protein interaction data were derived from publicly available interaction networks (e.g., BioPlex 3.0; https://bioplex.hms.harvard.edu).

Perturbational transcriptomes were obtained from the FRoGS repository^14^, originally sourced from the LINCS L1000 project (https://clue.io). PBMC3k single-cell data were accessed through the Scanpy dataset API, which retrieves the dataset from 10x Genomics (https://support.10xgenomics.com).

All custom code, including model training, embedding fusion, compound-target prediction, and visualization modules, is available at https://github.com/KidderLab/NEWT, along with environment files and reproducible workflows.

### COMPETING INTERESTS

The authors declare no conflict of interest.

### AUTHORS’ CONTRIBUTIONS

B.L.K. conceptualized the study, designed and performed the computational work, performed data analysis, generated the computational code, and prepared the manuscript.

## ACKNOWLEDGEMENTS

We also thank the Wayne State University High Performance Computing Grid (https://www.grid.wayne.edu/) for access to the computational resources that made this work possible.

## FUNDING

Karmanos Cancer Institute, Wayne State University; Karmanos Cancer Institute [P30 CA022453-Cancer Center Support Grant].

## Notes

### Competing Interest Statement

The authors have declared no competing interest.

## REFERENCES

1. Lamb J, Crawford ED, Peck D, Modell JW, Blat IC, Wrobel MJ, Lerner J, Brunet JP, Subramanian A, Ross KN, Reich M, Hieronymus H, Wei G, Armstrong SA, Haggarty SJ, Clemons PA, Wei R, Carr SA, Lander ES, Golub TR. The Connectivity Map: using gene-expression signatures to connect small molecules, genes, and disease. Science. 2006;313(5795):1929–35. doi: 10.1126/science.1132939. PubMed PMID: 17008526.

2. Subramanian A, Narayan R, Corsello SM, Peck DD, Natoli TE, Lu X, Gould J, Davis JF, Tubelli AA, Asiedu JK, Lahr DL, Hirschman JE, Liu Z, Donahue M, Julian B, Khan M, Wadden D, Smith IC, Lam D, Liberzon A, Toder C, Bagul M, Orzechowski M, Enache OM, Piccioni F, Johnson SA, Lyons NJ, Berger AH, Shamji AF, Brooks AN, Vrcic A, Flynn C, Rosains J, Takeda DY, Hu R, Davison D, Lamb J, Ardlie K, Hogstrom L, Greenside P, Gray NS, Clemons PA, Silver S, Wu X, Zhao WN, Read-Button W, Wu X, Haggarty SJ, Ronco LV, Boehm JS, Schreiber SL, Doench JG, Bittker JA, Root DE, Wong B, Golub TR. A Next Generation Connectivity Map: L1000 Platform and the First 1,000,000 Profiles. Cell. 2017;171(6):1437–52 e17. doi: 10.1016/j.cell.2017.10.049. PubMed PMID: 29195078; PMCID: PMC5990023.

3. Iorio F, Knijnenburg TA, Vis DJ, Bignell GR, Menden MP, Schubert M, Aben N, Goncalves E, Barthorpe S, Lightfoot H, Cokelaer T, Greninger P, van Dyk E, Chang H, de Silva H, Heyn H, Deng X, Egan RK, Liu Q, Mironenko T, Mitropoulos X, Richardson L, Wang J, Zhang T, Moran S, Sayols S, Soleimani M, Tamborero D, Lopez-Bigas N, Ross-Macdonald P, Esteller M, Gray NS, Haber DA, Stratton MR, Benes CH, Wessels LFA, Saez-Rodriguez J, McDermott U, Garnett MJ. A Landscape of Pharmacogenomic Interactions in Cancer. Cell. 2016;166(3):740–54. doi: 10.1016/j.cell.2016.06.017. PubMed PMID: 27397505; PMCID: PMC4967469.

4. DiMasi JA, Grabowski HG, Hansen RW. Innovation in the pharmaceutical industry: New estimates of R&D costs. J Health Econ. 2016;47:20–33. doi: 10.1016/j.jhealeco.2016.01.012. PubMed PMID: 26928437.

5. Barabasi AL, Gulbahce N, Loscalzo J. Network medicine: a network-based approach to human disease. Nat Rev Genet. 2011;12(1):56–68. doi: 10.1038/nrg2918. PubMed PMID: 21164525; PMCID: PMC3140052.

6. Guney E, Menche J, Vidal M, Barabasi AL. Network-based in silico drug efficacy screening. Nat Commun. 2016;7:10331. doi: 10.1038/ncomms10331. PubMed PMID: 26831545; PMCID: PMC4740350.

7. Zitnik M, Agrawal M, Leskovec J. Modeling polypharmacy side effects with graph convolutional networks. Bioinformatics. 2018;34(13):i457–i66. doi: 10.1093/bioinformatics/bty294. PubMed PMID: 29949996; PMCID: PMC6022705.

8. Hamilton WL, Ying R, Leskovec J. Inductive Representation Learning on Large Graphs. arXiv; 2018.

9. Zhao Y, Xing Y, Zhang Y, Wang Y, Wan M, Yi D, Wu C, Li S, Xu H, Zhang H, Liu Z, Zhou G, Li M, Wang X, Chen Z, Li R, Wu L, Zhao D, Zan P, He S, Bo X. Evidential deep learning-based drug-target interaction prediction. Nat Commun. 2025;16(1):6915. doi: 10.1038/s41467-025-62235-6. PubMed PMID: 40715097; PMCID: PMC12297561.

10. Vamathevan J, Clark D, Czodrowski P, Dunham I, Ferran E, Lee G, Li B, Madabhushi A, Shah P, Spitzer M, Zhao S. Applications of machine learning in drug discovery and development. Nat Rev Drug Discov. 2019;18(6):463–77. doi: 10.1038/s41573-019-0024-5. PubMed PMID: 30976107; PMCID: PMC6552674.

11. Du J, Jia P, Dai Y, Tao C, Zhao Z, Zhi D. Gene2vec: distributed representation of genes based on co-expression. BMC Genomics. 2019;20(Suppl 1):82. doi: 10.1186/s12864-018-5370-x. PubMed PMID: 30712510; PMCID: PMC6360648.

12. Smaili FZ, Gao X, Hoehndorf R. OPA2Vec: combining formal and informal content of biomedical ontologies to improve similarity-based prediction. Bioinformatics. 2019;35(12):2133–40. doi: 10.1093/bioinformatics/bty933. PubMed PMID: 30407490.

13. Wang S, Cho H, Zhai C, Berger B, Peng J. Exploiting ontology graph for predicting sparsely annotated gene function. Bioinformatics. 2015;31(12):i357–64. doi: 10.1093/bioinformatics/btv260. PubMed PMID: 26072504; PMCID: PMC4542782.

14. Chen H, King FJ, Zhou B, Wang Y, Canedy CJ, Hayashi J, Zhong Y, Chang MW, Pache L, Wong JL, Jia Y, Joslin J, Jiang T, Benner C, Chanda SK, Zhou Y. Drug target prediction through deep learning functional representation of gene signatures. Nat Commun. 2024;15(1):1853. doi: 10.1038/s41467-024-46089-y. PubMed PMID: 38424040; PMCID: PMC10904399.

15. Svensson V, Vento-Tormo R, Teichmann SA. Exponential scaling of single-cell RNA-seq in the past decade. Nat Protoc. 2018;13(4):599–604. doi: 10.1038/nprot.2017.149. PubMed PMID: 29494575.

16. Stuart T, Satija R. Integrative single-cell analysis. Nat Rev Genet. 2019;20(5):257–72. doi: 10.1038/s41576-019-0093-7. PubMed PMID: 30696980.

17. Kelsey G, Stegle O, Reik W. Single-cell epigenomics: Recording the past and predicting the future. Science. 2017;358(6359):69–75. doi: 10.1126/science.aan6826. PubMed PMID: 28983045.

18. Ashburner M, Ball CA, Blake JA, Botstein D, Butler H, Cherry JM, Davis AP, Dolinski K, Dwight SS, Eppig JT, Harris MA, Hill DP, Issel-Tarver L, Kasarskis A, Lewis S, Matese JC, Richardson JE, Ringwald M, Rubin GM, Sherlock G. Gene ontology: tool for the unification of biology. The Gene Ontology Consortium. Nat Genet. 2000;25(1):25–9. doi: 10.1038/75556. PubMed PMID: 10802651; PMCID: PMC3037419.

19. Lachmann A, Torre D, Keenan AB, Jagodnik KM, Lee HJ, Wang L, Silverstein MC, Ma’ayan A. Massive mining of publicly available RNA-seq data from human and mouse. Nat Commun. 2018;9(1):1366. doi: 10.1038/s41467-018-03751-6. PubMed PMID: 29636450; PMCID: PMC5893633.

20. Liberzon A, Birger C, Thorvaldsdottir H, Ghandi M, Mesirov JP, Tamayo P. The Molecular Signatures Database (MSigDB) hallmark gene set collection. Cell Syst. 2015;1(6):417–25. doi: 10.1016/j.cels.2015.12.004. PubMed PMID: 26771021; PMCID: PMC4707969.

21. Cahan P, Li H, Morris SA, Lummertz da Rocha E, Daley GQ, Collins JJ. CellNet: network biology applied to stem cell engineering. Cell. 2014;158(4):903–15. doi: 10.1016/j.cell.2014.07.020. PubMed PMID: 25126793; PMCID: PMC4233680.

22. Garcia-Alonso L, Holland CH, Ibrahim MM, Turei D, Saez-Rodriguez J. Benchmark and integration of resources for the estimation of human transcription factor activities. Genome Res. 2019;29(8):1363–75. doi: 10.1101/gr.240663.118. PubMed PMID: 31340985; PMCID: PMC6673718.

23. Muller-Dott S, Tsirvouli E, Vazquez M, Ramirez Flores RO, Badia IMP, Fallegger R, Turei D, Laegreid A, Saez-Rodriguez J. Expanding the coverage of regulons from high-confidence prior knowledge for accurate estimation of transcription factor activities. Nucleic Acids Res. 2023;51(20):10934–49. doi: 10.1093/nar/gkad841. PubMed PMID: 37843125; PMCID: PMC10639077.

24. Luck K, Kim DK, Lambourne L, Spirohn K, Begg BE, Bian W, Brignall R, Cafarelli T, Campos-Laborie FJ, Charloteaux B, Choi D, Cote AG, Daley M, Deimling S, Desbuleux A, Dricot A, Gebbia M, Hardy MF, Kishore N, Knapp JJ, Kovacs IA, Lemmens I, Mee MW, Mellor JC, Pollis C, Pons C, Richardson AD, Schlabach S, Teeking B, Yadav A, Babor M, Balcha D, Basha O, Bowman-Colin C, Chin SF, Choi SG, Colabella C, Coppin G, D’Amata C, De Ridder D, De Rouck S, Duran-Frigola M, Ennajdaoui H, Goebels F, Goehring L, Gopal A, Haddad G, Hatchi E, Helmy M, Jacob Y, Kassa Y, Landini S, Li R, van Lieshout N, MacWilliams A, Markey D, Paulson JN, Rangarajan S, Rasla J, Rayhan A, Rolland T, San-Miguel A, Shen Y, Sheykhkarimli D, Sheynkman GM, Simonovsky E, Tasan M, Tejeda A, Tropepe V, Twizere JC, Wang Y, Weatheritt RJ, Weile J, Xia Y, Yang X, Yeger-Lotem E, Zhong Q, Aloy P, Bader GD, De Las Rivas J, Gaudet S, Hao T, Rak J, Tavernier J, Hill DE, Vidal M, Roth FP, Calderwood MA. A reference map of the human binary protein interactome. Nature. 2020;580(7803):402–8. doi: 10.1038/s41586-020-2188-x. PubMed PMID: 32296183; PMCID: PMC7169983.

25. Zheng GX, Terry JM, Belgrader P, Ryvkin P, Bent ZW, Wilson R, Ziraldo SB, Wheeler TD, McDermott GP, Zhu J, Gregory MT, Shuga J, Montesclaros L, Underwood JG, Masquelier DA, Nishimura SY, Schnall-Levin M, Wyatt PW, Hindson CM, Bharadwaj R, Wong A, Ness KD, Beppu LW, Deeg HJ, McFarland C, Loeb KR, Valente WJ, Ericson NG, Stevens EA, Radich JP, Mikkelsen TS, Hindson BJ, Bielas JH. Massively parallel digital transcriptional profiling of single cells. Nat Commun. 2017;8:14049. doi: 10.1038/ncomms14049. PubMed PMID: 28091601; PMCID: PMC5241818 L.M., D.A.M., S.Y.N., M.S.L., P.W.W., C.M.H., R.B., A.W., K.D.N., T.S.M. and B.J.H. are employees of 10x Genomics.

26. Gong P, Cheng L, Zhang Z, Meng A, Li E, Chen J, Zhang L. Multi-omics integration method based on attention deep learning network for biomedical data classification. Comput Methods Programs Biomed. 2023;231:107377. doi: 10.1016/j.cmpb.2023.107377. PubMed PMID: 36739624.

27. Sun Q, Cheng L, Meng A, Ge S, Chen J, Zhang L, Gong P. SADLN: Self-attention based deep learning network of integrating multi-omics data for cancer subtype recognition. Front Genet. 2022;13:1032768. doi: 10.3389/fgene.2022.1032768. PubMed PMID: 36685873; PMCID: PMC9846505.

28. Vaswani A, Shazeer N, Parmar N, Uszkoreit J, Jones L, Gomez AN, Kaiser L, Polosukhin I. Attention Is All You Need. arXiv preprint arXiv:1706.03762; 2017.

29. Wolf FA, Angerer P, Theis FJ. SCANPY: large-scale single-cell gene expression data analysis. Genome Biol. 2018;19(1):15. doi: 10.1186/s13059-017-1382-0. PubMed PMID: 29409532; PMCID: PMC5802054.

30. Szklarczyk D, Kirsch R, Koutrouli M, Nastou K, Mehryary F, Hachilif R, Gable AL, Fang T, Doncheva NT, Pyysalo S, Bork P, Jensen LJ, von Mering C. The STRING database in 2023: protein-protein association networks and functional enrichment analyses for any sequenced genome of interest. Nucleic Acids Res. 2023;51(D1):D638–D46. doi: 10.1093/nar/gkac1000. PubMed PMID: 36370105; PMCID: PMC9825434.

31. Wishart DS, Feunang YD, Guo AC, Lo EJ, Marcu A, Grant JR, Sajed T, Johnson D, Li C, Sayeeda Z, Assempour N, Iynkkaran I, Liu Y, Maciejewski A, Gale N, Wilson A, Chin L, Cummings R, Le D, Pon A, Knox C, Wilson M. DrugBank 5.0: a major update to the DrugBank database for 2018. Nucleic Acids Res. 2018;46(D1):D1074–D82. doi: 10.1093/nar/gkx1037. PubMed PMID: 29126136; PMCID: PMC5753335.

32. Zdrazil B, Felix E, Hunter F, Manners EJ, Blackshaw J, Corbett S, de Veij M, Ioannidis H, Lopez DM, Mosquera JF, Magarinos MP, Bosc N, Arcila R, Kiziloren T, Gaulton A, Bento AP, Adasme MF, Monecke P, Landrum GA, Leach AR. The ChEMBL Database in 2023: a drug discovery platform spanning multiple bioactivity data types and time periods. Nucleic Acids Res. 2024;52(D1):D1180–D92. doi: 10.1093/nar/gkad1004. PubMed PMID: 37933841; PMCID: PMC10767899.

33. Shannon P, Markiel A, Ozier O, Baliga NS, Wang JT, Ramage D, Amin N, Schwikowski B, Ideker T. Cytoscape: a software environment for integrated models of biomolecular interaction networks. Genome Res. 2003;13(11):2498–504. doi: 10.1101/gr.1239303. PubMed PMID: 14597658; PMCID: PMC403769.

