## Supplemental Information for "Multimodal gene embeddings for drug-target prediction and lineage reconstruction"

Benjamin L. Kidder[1-2*]

[1]Department of Oncology, Wayne State University School of Medicine, Detroit, MI, USA

[2]Karmanos Cancer Institute, Wayne State University School of Medicine, Detroit, MI, USA

Running title: Multimodal embeddings for target and lineage mapping .

*Correspondence:

Benjamin L. Kidder

### SUPPLEMENTARY FIGURES

**Figure S1. Comparison of intermediate multimodal-only embedding with final joint model.**

(A) UMAP projection of the intermediate multimodal embedding showing lineage organization. (B) Dotplot and (C) heatmap visualizations of canonical immune marker expression across clusters.

**Figure S2. Stability of modular architecture across normalization schemes.**

(A–D) Comparative heatmaps showing lineage marker expression across embeddings generated using different normalization methods: (A) FRoGS-default, (B) intermediate NEWT multimodal, (C) PCA baseline, and (D) NEWT joint model.

**Figure S3. Gene-level expression trajectories across multimodal embeddings.**

(A–D) Track plots showing expression continuity for representative lineage markers across clusters in (A) FRoGS-default, (B) intermediate NEWT multimodal, (C) PCA, and (D) NEWT joint embeddings.

**Figure S4. Expression distributions of canonical immune lineage markers across embeddings.**

(A–D) Heatmaps showing mean expression of representative lineage markers across clusters in (A) FRoGS-default, (B) intermediate NEWT multimodal, (C) PCA, and (D) NEWT joint embeddings.

**Figure S5. Expression distributions of B-cell markers across embeddings.**

(A–B) FRoGS-default embeddings showing broad, multimodal expression distributions for CD79A and MS4A1 across clusters. (C–D) Intermediate NEWT multimodal embeddings demonstrating partial sharpening of lineage specificity. (E–F) PCA-based embeddings displaying diffuse and overlapping expression profiles. (G–H) NEWT joint embeddings exhibiting unimodal, lineage-restricted expression confined to B-cell clusters, confirming precise marker localization.

**Figure S6. Quality-control metrics across embeddings.**

(A–B) FRoGS-default embeddings showing global distributions of the number of detected genes (n_genes) and mitochondrial read fraction (percent_mito) across clusters. (C–D) Intermediate NEWT multimodal embeddings demonstrating comparable distributions across clusters. (E–F) PCA-based embeddings displaying similar n_genes and percent_mito variability. (G–H) NEWT joint embeddings showing consistent profiles with no systematic bias across clusters.

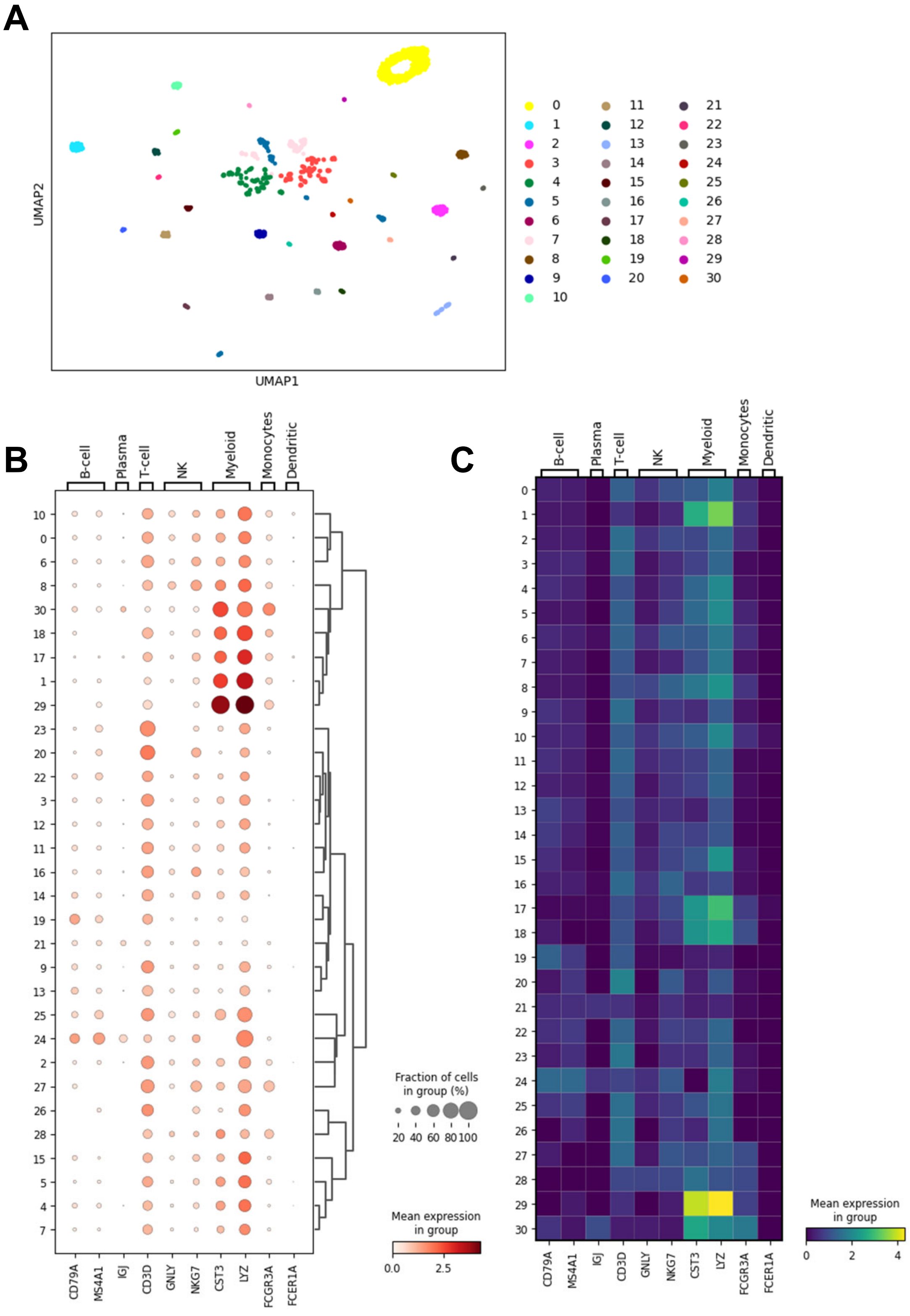
A
UMAP1
UMAP2
0 1 2 3 4 5 6 7 8 9 10 11 12 13 14 15 16 17 18 19 20 21 22 23 24 25 26 27 28 29 30
B
B-cell
Plasma
T-cell
NK
Myeloid
Monocytes
Dendritic
10 0 6 8 30 18 17 1 29 23 20 22 3 12 11 16 14 19 21 9 13 25 24 2 27 26 28 15 5 4 7
CD79A MS4A1 IGJ CD3D GNLY NKG7 CST3 LYZ FCGR3A FCER1A
Fraction of cells in group (%)
20 40 60 80 100
Mean expression in group
0.0 2.5
C
0 1 2 3 4 5 6 7 8 9 10 11 12 13 14 15 16 17 18 19 20 21 22 23 24 25 26 27 28 29 30
Mean expression in group
0 2 4

**Figure S1**

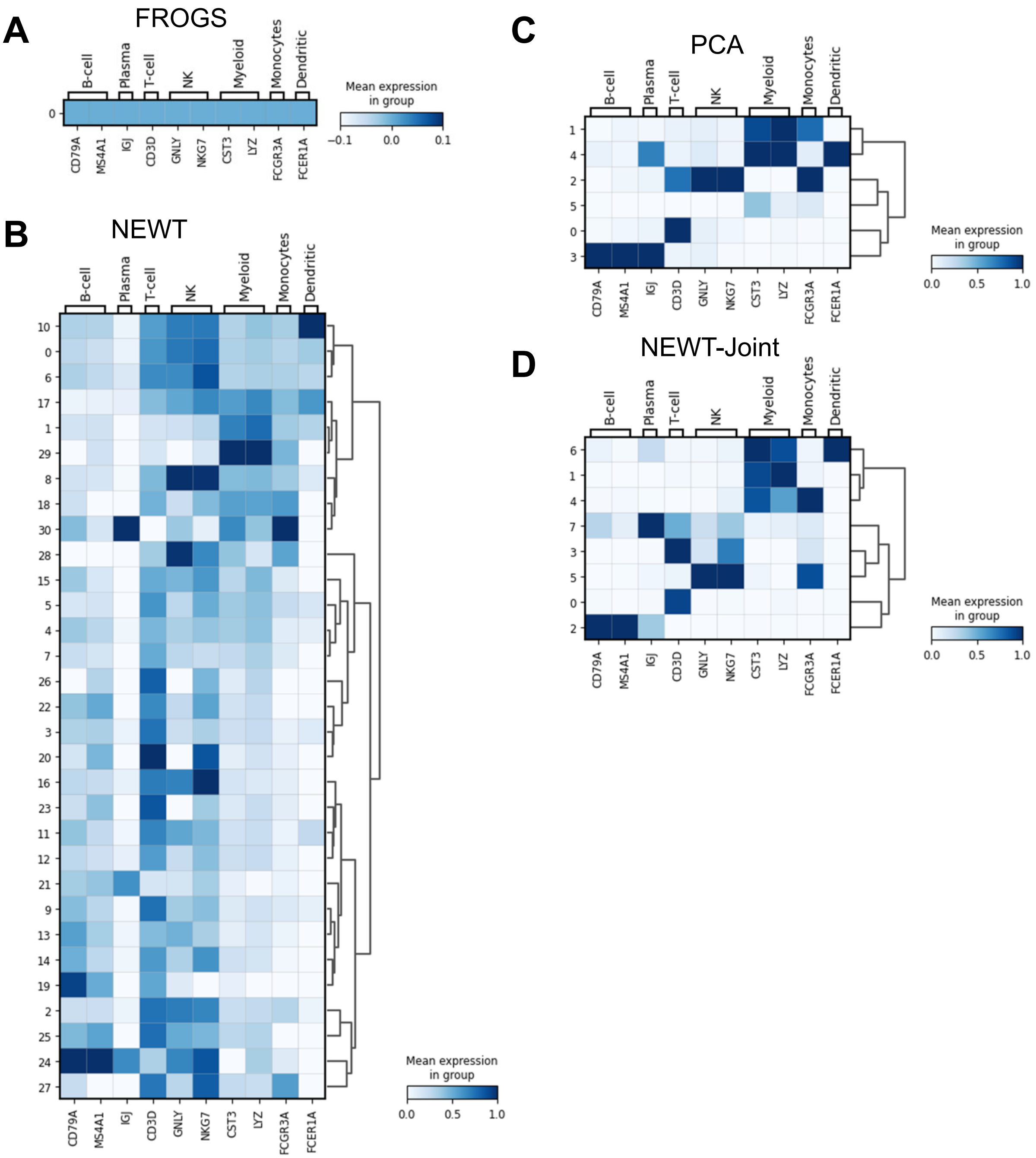
A
FROGS
B-cell
Plasma
T-cell
NK
Myeloid
Monocytes
Dendritic
0
CD79A
MS4A1
IGJ
CD3D
GNLY
NKG7
CST3
LYZ
FCGR3A
FCER1A
Mean expression in group
−0.1
0.0
0.1
B
NEWT
10
0
6
17
1
29
8
18
30
28
15
5
4
7
26
22
3
20
16
23
11
12
21
9
13
14
19
2
25
24
27
Mean expression in group
0.0
0.5
1.0
C
PCA
1
4
2
5
0
3
D
NEWT-Joint
6
1
4
7
3
5
0
2

Figure S2

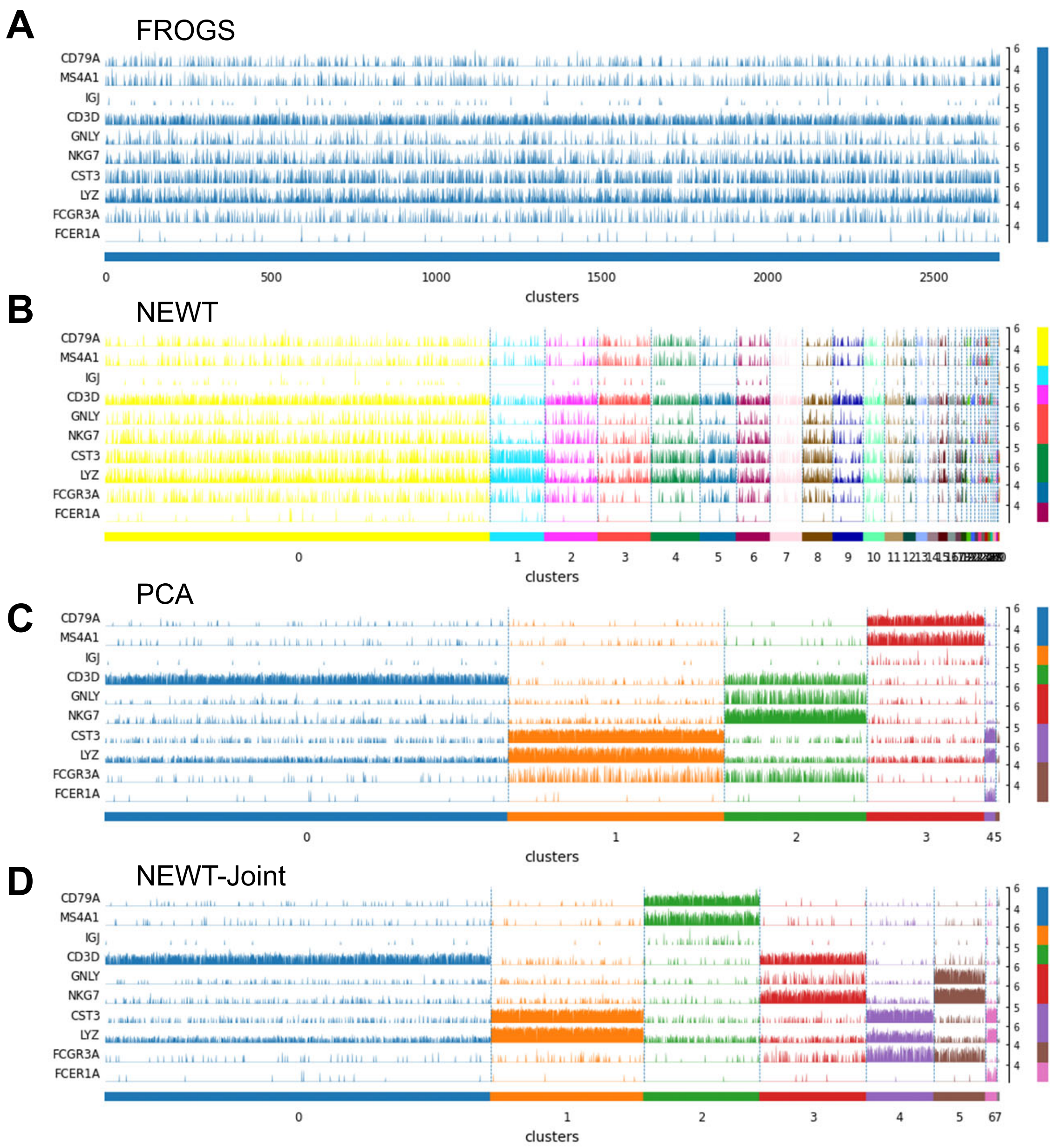
A
FROGS
CD79A
MS4A1
IGJ
CD3D
GNLY
NKG7
CST3
LYZ
FCGR3A
FCER1A
0
500
1000
1500
2000
2500
clusters
B
NEWT
clusters
C
PCA
clusters
D
NEWT-Joint
clusters

**Figure S3**

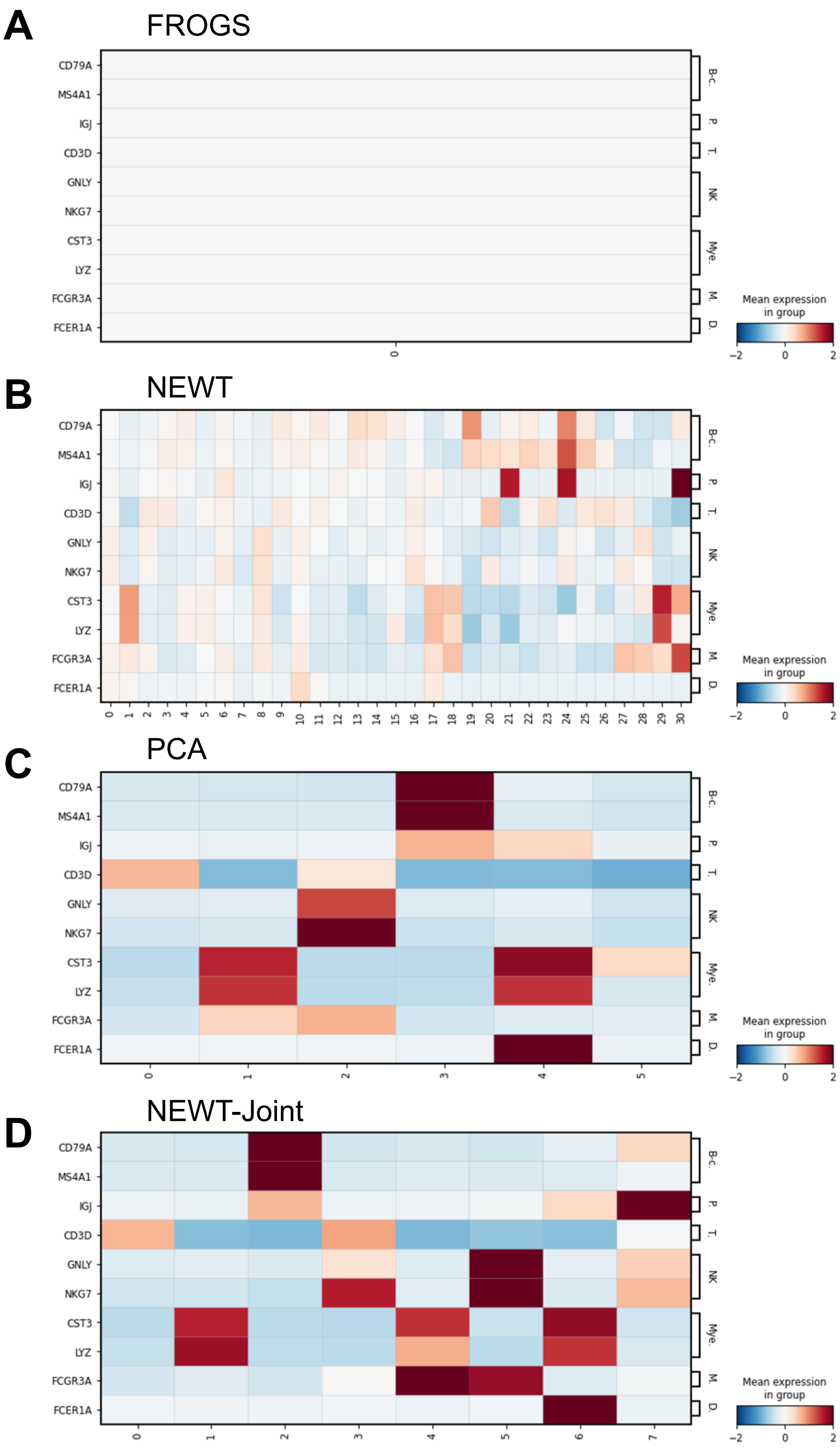
A
FROGS
B
NEWT
C
PCA
D
NEWT-Joint
CD79A
MS4A1
IGJ
CD3D
GNLY
NKG7
CST3
LYZ
FCGR3A
FCER1A
B-c.
P.
T.
NK
Mye.
M.
D.
Mean expression in group
−2
0
2

**Figure S4**

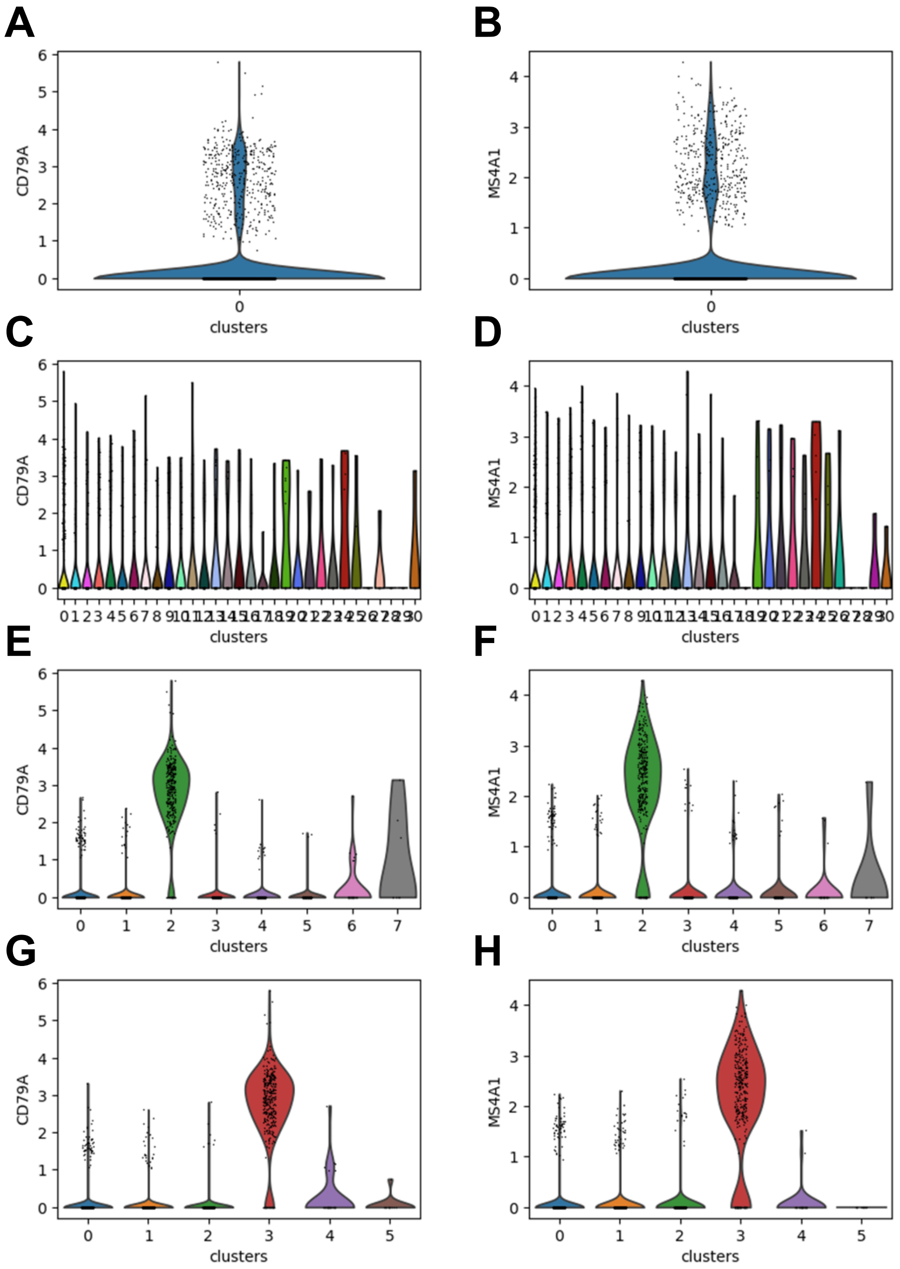


**Figure S5**

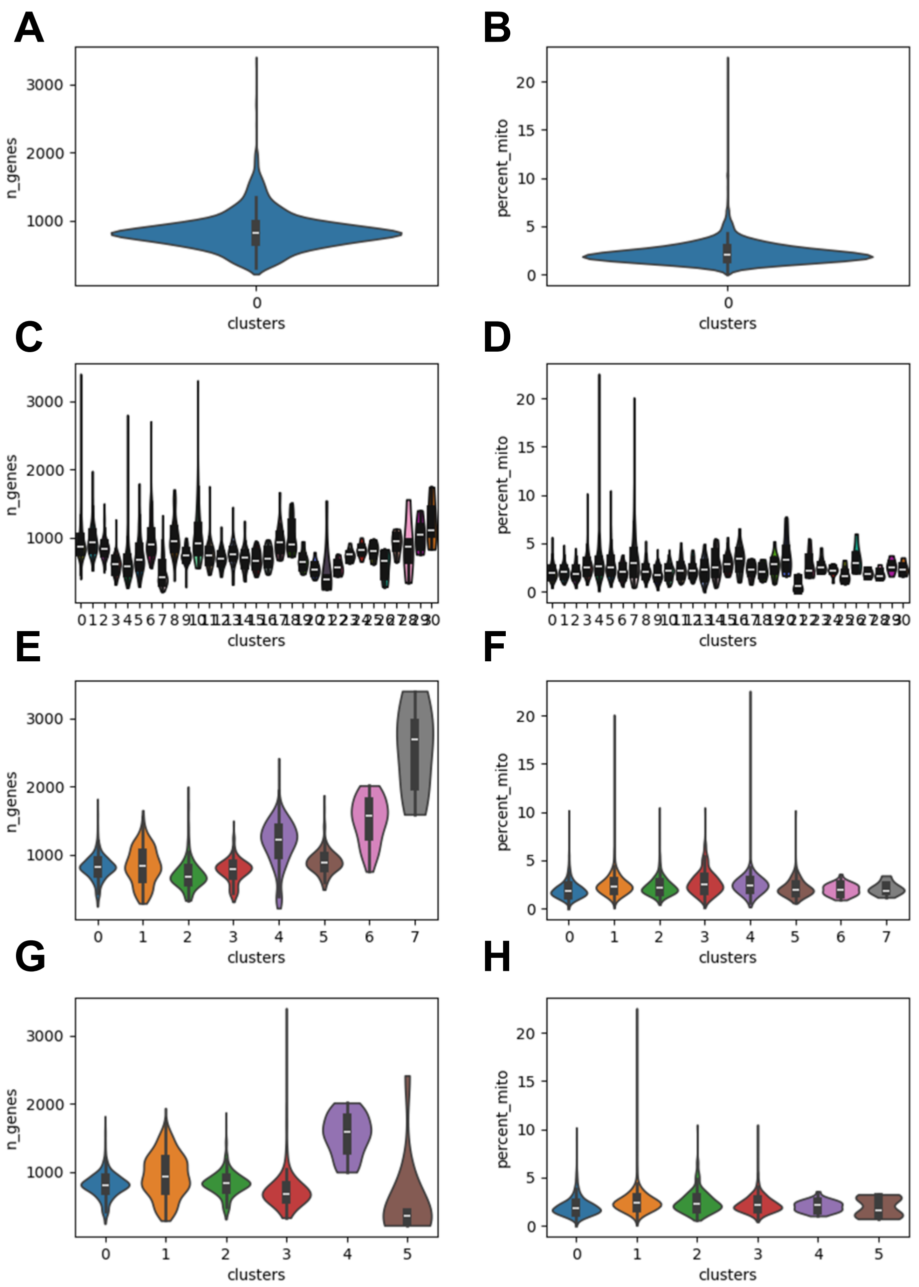
A
B
C
D
E
F
G
H
n_genes
percent_mito
clusters
3000
2000
1000
20
15
10
5
0
0
1
2
3
4
5
6
7

**Figure S6**
